# Acute stress enhances associative learning via dopamine signaling in the ventral lateral striatum

**DOI:** 10.1101/2019.12.18.881417

**Authors:** Claire E. Stelly, Sean C. Tritley, Yousef Rafati, Matthew J. Wanat

## Abstract

Acute stress transiently increases vigilance, whereby enhancing the detection of salient stimuli. This increased perceptual sensitivity is thought to promote associating rewarding outcomes with relevant cues. The mesolimbic dopamine system is critical for learning cue-reward associations. Dopamine levels in the ventral striatum are elevated following exposure to stress. Together, this suggests the mesolimbic dopamine system could mediate the influence of acute stress on cue-reward learning. To address this possibility, we examined how a single stressful experience influenced learning in an appetitive Pavlovian conditioning task. Male rats underwent an episode of restraint prior to the first conditioning session. This acute stress treatment augmented conditioned responding in subsequent sessions. Voltammetry recordings of mesolimbic dopamine levels demonstrated that acute stress selectively increased reward-evoked dopamine release in the ventral lateral striatum (VLS), but not in the ventral medial striatum (VMS). Antagonizing dopamine receptors in the VLS blocked the stress-induced enhancement of conditioned responding. Collectively, these findings illustrate that stress engages dopamine signaling in the VLS to facilitate appetitive learning.

## Introduction

Acute stress triggers a transient state of increased vigilance. This heightened awareness of one’s surroundings reflects activation of the ‘salience network’, a large-scale brain network for detecting and attending to stimuli that are potentially harmful or beneficial^1-5^. Increased stimulus salience is theorized to facilitate associative learning^6,7^. As stress increases salience, associative learning should be enhanced accordingly. Consistent with this idea, stress promotes conditioned responding to aversive cues^8-12^. While stress facilitates learning to associate contextual cues with drug rewards^13,14^, it is unclear if acute stress additionally enhances conditioning with natural rewards.

Phasic dopamine release in the ventral striatum is essential for learning to associate cues with rewarding outcomes^15-19^. The mesolimbic dopamine system is also sensitive to stress, as dopamine levels in the ventral striatum are modulated during and after exposure to stressors^20-25^. However, it is not known if acute stress regulates phasic dopamine release to impact associative learning.

To address this question, male rats were exposed to a single episode of restraint stress prior to training on a Pavlovian conditioning task using food rewards. We monitored dopamine release in the ventral medial and ventral lateral striatum throughout training to determine if stress altered the dopamine response to rewards or their predictors. Additionally, we performed local pharmacological manipulations to establish if stress-induced behavioral changes required dopamine transmission.

## Methods

### Subjects and surgery

The University of Texas at San Antonio Institutional Animal Care and Use Committee approved all procedures. Male CD IGS Sprague Dawley rats (Charles River Laboratories, RRID:RGD 734476) were pair-housed upon arrival, allowed *ad libitum* access to water and chow, and maintained on a 12 h light/dark cycle. Voltammetry electrodes were surgically implanted under isoflurane anesthesia in rats weighing 300 – 400 g. Carbon fiber electrodes were placed bilaterally targeting the VMS or VLS (relative to bregma: 1.3 mm anterior; ± 1.3 mm lateral; 7.0 mm ventral or 1.3 mm anterior; ± 2.7 mm lateral; 7.3 mm ventral, respectively), along with an Ag/AgCl reference electrode placed under the skull. Bilateral stainless steel guide cannulae (InVivo One) were implanted 1 mm dorsal to the VLS. Following surgery, rats were single-housed for the duration of the experiment and allowed to recover for 1-3 weeks before behavioral procedures. Electrode and cannula placements are depicted in Fig. 1. The microinjection area is based on the spread of an equivalent volume of Evans Blue dye.

**Figure 1.**
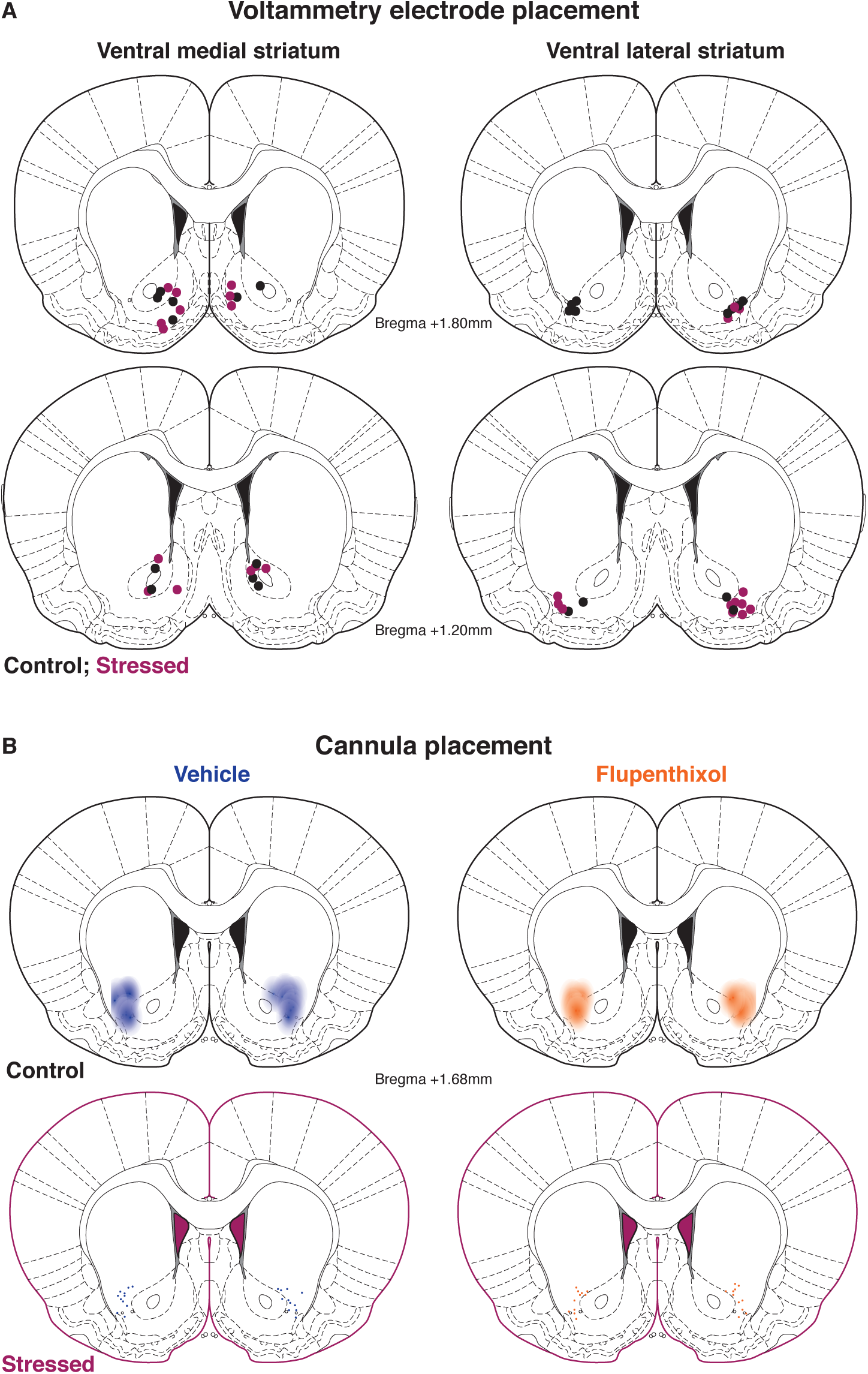
Voltammetry electrode and cannula placement. **A**, Histologically verified locations of voltammetry electrodes in control rats (black circles) and stressed rats (magenta circles). We used the lateral edge of the anterior commissure as the boundary of the ventral medial (left) and ventral lateral (right) striatum. **B**, Histologically verified locations of microinjector tips and approximate infusion area of vehicle (left, blue) and flupenthixol (right, orange) in control rats (above, black border) and stressed rats (below, magenta border).

### Behavioral procedures

At ≥ 7 days post-surgery, rats were placed on mild dietary restriction to 90% of their free feeding weight, allowing for a weekly increase of 1.5%. Animals were handled regularly before behavioral testing commenced. All behavioral sessions occurred during the light cycle in operant boxes (Med Associates) with a grid floor, a house light, a recessed food tray equipped with an infrared beam-break detector, and a white noise generator. To familiarize the animals with the operant chamber and food retrieval from the tray, rats first received 1-2 magazine training sessions in which 20 unsignaled food pellets (45 mg, BioServ) were delivered at a 90 s variable interval. Rats then underwent 10 Pavlovian reward conditioning sessions comprised of 50 trials each. Trials consisted of a 5 s white noise conditioned stimulus (CS) presentation terminating with the delivery of a single food pellet unconditioned stimulus (US) and 4.5 s illumination of the tray light. Trials were separated by a 55 ± 15 s intertrial interval. We monitored head entries into the food tray across training sessions. Conditioned responding was quantified as the change in the rate of head entries during the 5 s CS relative to the 5 s preceding the CS delivery^26^. Response latency was calculated as the interval from CS onset to the first head entry during the CS. To assay response vigor, we calculated the head entry rate during the interval from the first entry to the end of the CS. We then took the difference between this adjusted response rate relative to the head entry rate in the 5 s preceding the CS delivery.

### Restraint stress

In a novel room, rats were either introduced to a clean, empty cage (control) or confined in a clear acrylic tail vein restrainer (Braintree Scientific) for 20-30 min (stress). Rats were then transferred to a clean recovery cage in the familiar testing area for 5 min. Following recovery, rats were connected to the voltammetric amplifier in the operant chamber and electrodes were cycled for 15 min prior to Pavlovian training sessions, for a total interval of 20 min from the end of stress/control procedure to the start of training. An additional group of animals were returned to their home cages for 1o0 min after recovery, allowing for a 2 hr interval from the end of stress/control procedure to the start of training.

### Pharmacology

Flupenthixol dihydrochloride (Tocris) was dissolved in sterile 0.9% NaCl. Rats received bilateral 0.5 µl microinjections of flupenthixol (10 µg/side) or vehicle into the ventrolateral striatum at 0.25 µl/min. The injectors were removed 1 minute after the infusion ended. Behavioral sessions commenced 30 min after the intra-VLS microinjections^27^.

### Voltammetry recordings and analysis

Indwelling carbon fiber microelectrodes were connected to a head-mounted amplifier to monitor dopamine release in behaving rats using fast-scan cyclic voltammetry as described previously^26,28-30^. During voltammetric scans, the potential applied to the carbon fiber was ramped in a triangular waveform from −0.4 V (vs. Ag/ AgCl) to +1.3 V and back at a rate of 400 V/s. Scans occurred at 10 Hz with the electrode potential held at −0.4 V between scans. Dopamine was chemically verified by obtaining high correlation of the cyclic voltammogram during a reward-related event with that of a dopamine standard (correlation coefficient *r*^2^ ≥ 0.75 by linear regression). Voltammetry data for a session were excluded from analysis if the detected voltammetry signal did not satisfy the chemical verification criteria, as in prior studies^26,29,30^. Dopamine was isolated from the voltammetry signal using chemometric analysis^31^ with a standard training set accounting for dopamine, pH, and drift. The background for voltammetry recording analysis was set at 0.5 s before the CS onset. CS-evoked dopamine release was quantified as the mean dopamine level during the 5 s CS relative to the 5 s prior to the CS delivery^26^. US-evoked dopamine was quantified as the peak dopamine level during the 2.5 s following US delivery relative to the 0.5 s preceding the US delivery. Trials were excluded if chemometric analysis failed to identify dopamine on > 25% of the data points. The change in dopamine concentration was estimated based on the average post-implantation electrode sensitivity (34 nA/µM)^28^.

### Experimental design and statistical analysis

Rats were assigned to stress or control groups in an unbiased manner. We performed all statistical analyses in Graphpad Prism 8. All data are plotted as mean ± SEM. A mixed-effects model fit (restricted maximum likelihood method) was used to analyze effects on behavioral measures and dopamine responses. Student’s unpaired t-test with Welch’s correction was used to compare dopamine responses between dorsal and ventral VMS. The significance level was set to α = 0.05 for all tests.

### Histology

Rats with were anesthetized, electrically lesioned via the voltammetry electrodes, and perfused intracardially with 4% paraformaldehyde. Brains were extracted and post-fixed in the paraformaldehyde solution for a minimum of 24 hrs, then were transferred to 15 % and 30 % sucrose in phosphate-buffered saline. Tissue was cryosectioned and stained with cresyl violet. Implant locations were mapped to a standardized rat brain atlas^32^. The VMS and VLS were delineated by the anatomical boundary formed by the lateral edge of the anterior commissure.

## Results

### A single stress exposure enhances conditioned responding to reward-predictive stimuli

We examined how a single episode of restraint stress affected the acquisition of conditioned behavioral responses to a reward-predictive cue. As a control, a separate group of rats was exposed to a clean, empty cage for an equivalent period of time. Rats underwent the stress or control treatment 20 min prior to the first Pavlovian conditioning session. Training continued for 9 additional daily sessions without any further stress experience (Fig. 2A). Each session consisted of 50 presentations of a 5 s audio CS that terminated with the delivery of a single food pellet US (Fig. 2B). Conditioned responding was elevated in stressed rats relative to controls (treatment effect *F*_(1, 35)_=8.22, *p=*0.007; *n*=16 control, 21 stress; Fig. 2C). Stress did not alter the time to approach to the food tray, as the latency from the CS onset to the first tray entry did not differ between groups (treatment effect *F*_(1, 35)_=0.80, *p=*0.38; Fig. 2D). The number of tray entries during the intertrial interval was unaffected by stress exposure, indicating no change in overall activity (treatment effect *F*_(1, 35)_=1.03, *p=*0.32; Fig. 2E). Together, these results demonstrate that stress selectively increases conditioned responses towards a reward-predictive cue.

**Figure 2.**
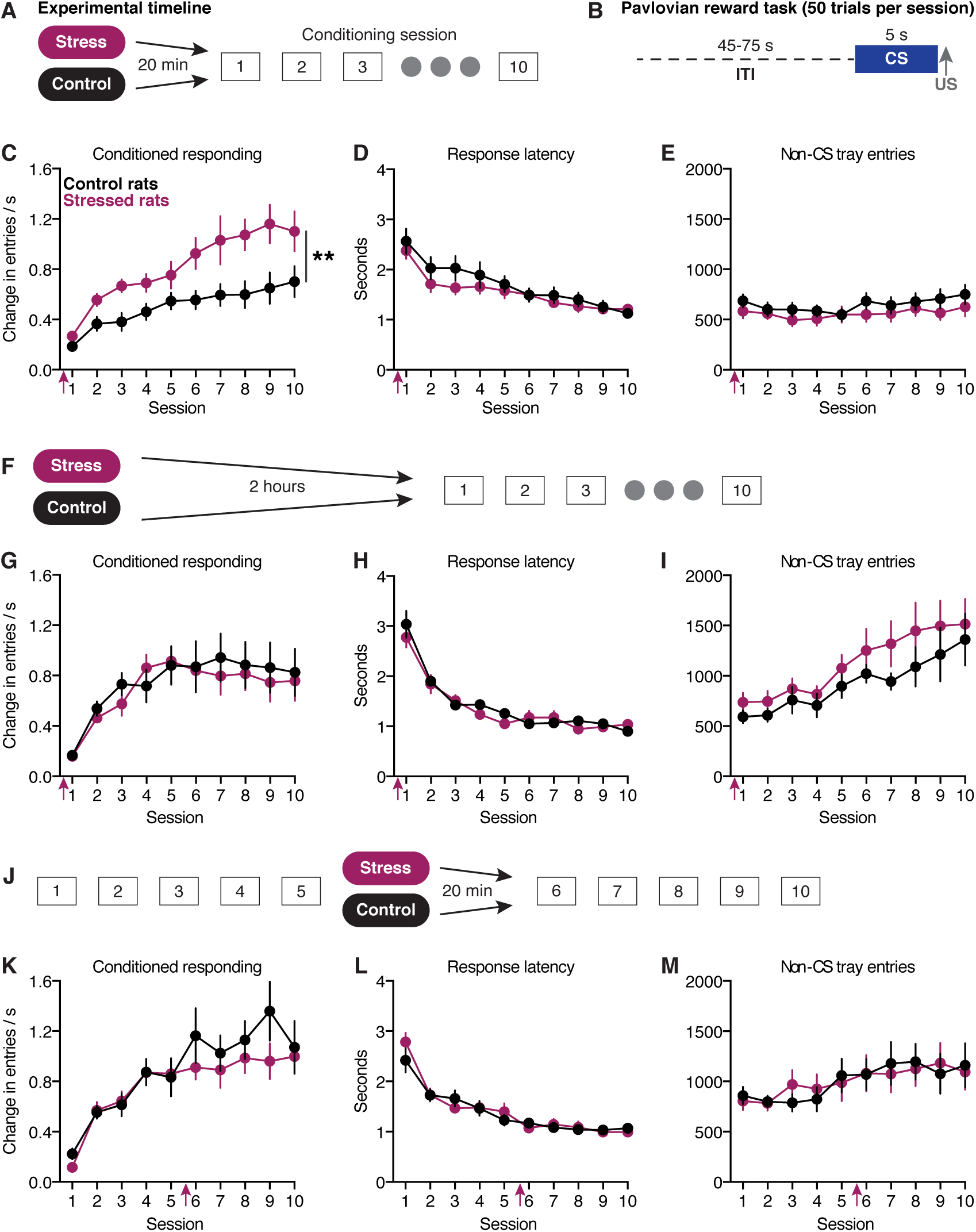
A single stress experience enhances subsequent Pavlovian conditioning. **A**, Training paradigm. Animals are stressed once, 20 min prior to the first conditioning session. **B**, Task structure. **C**, Elevated conditioned responding to the reward-predictive CS in rats stressed before the first training session. Magenta arrows denote restraint stress/control procedure **D**, Latency to head entry. **E**, Non-CS tray entries. **F**, Training paradigm with a 2 hr delay between the stress/control treatment and the start of conditioning. **G**, Conditioned responding is not increased when training begins 2 hrs after the stressor. **H**, Latency to head entry. **I**, Non-CS tray entries. **J**, Training paradigm with stress/control treatment occurring 20 min prior to the sixth conditioning session. **K**, Conditioned responding is not increased when stress experience occurs after acquisition of the task. **L**, Latency to head entry. **M**, Non-CS tray entries. \*\**p* < .01

Stressful experience produces physiological effects ranging from minutes to hours^33^. To determine the temporal window in which acute stress impacts Pavlovian reward learning, we increased the interval between the stressor and the conditioning session to 2 hrs (Fig. 2F). Conditioned responding did not differ between stressed and control rats (treatment effect *F*_(1, 17)_=0.13, *p*=0.72; *n*=9 control, 10 stress; Fig. 2G). Additionally, there was no difference in the response latency (treatment effect *F*_(1, 17)_=0.28, *p*=0.61; Fig.2H) or non-CS tray entries (treatment effect *F*_(1, 17)_=1.20, *p*=0.29; Fig. 2I). These findings demonstrate that the stress exposure and the training experience must occur in close temporal proximity for stress to affect learning.

We next examined if stress exposure similarly facilitated conditioned responding in well-trained animals. Rats were trained for 5 Pavlovian conditioning sessions before undergoing a single stress exposure 20 min prior to the sixth session (Fig. 2J). Acute stress exposure in well-trained animals did not impact subsequent conditioned responding (treatment effect *F*_(1, 2o)_=1.07, *p*=0.31; *n*=10 control, 12 stress; Fig. 2K), response latency (treatment effect *F*_(1, 20)_=0.05, *p*=0.82; Fig. 2L) or non-CS tray entries (treatment effect *F*_(1, 20)_=0.01, *p*=0.93; Fig. 2M). These results illustrate that acute stress does not influence the expression of a previously acquired conditioned response.

### Stress selectively enhances reward-evoked dopamine release in the ventral lateral striatum

Dopamine transmission in the ventral striatum is required for the acquisition of conditioned responding^17^. Furthermore, increasing ventral striatal phasic dopamine release is sufficient to confer conditioned motivational properties to neutral stimuli^19^. The enhanced conditioned responding observed after acute stress could therefore reflect stress-induced augmentation of ventral striatal dopamine signals. To address this possibility, we performed voltammetry recordings of dopamine release in the ventral striatum across Pavlovian conditioning sessions. We analyzed dopamine release during the first five sessions, as conditioned responding was insensitive to the stress manipulation after this point (Fig 2J-M).

We first examined dopamine signaling in the ventral medial striatum (VMS) given the involvement of the VMS in reward-related behaviors^34,35^. Consistent with prior studies^36,37^, dopamine release in the VMS was time-locked to both the CS and the US (Fig. 3B-C). The CS dopamine response did not differ between stressed and control rats (treatment effect *F*_(1, 20)=_0.64, *p*=0.43; *n*=9 control, 13 stress; Fig. 3D). Dopamine release to the US decayed with training (session effect *F*_(2.5, 43.9)_=8.02, *p*=.0005; Fig 3E), but was unaffected by stress exposure (treatment effect *F*_(1, 20)_=2.22, *p*=0.15). Collectively, these results indicate that acute stress prior to the first conditioning session did not influence the VMS dopamine response to rewards or reward-predictive cues.

**Figure 3.**
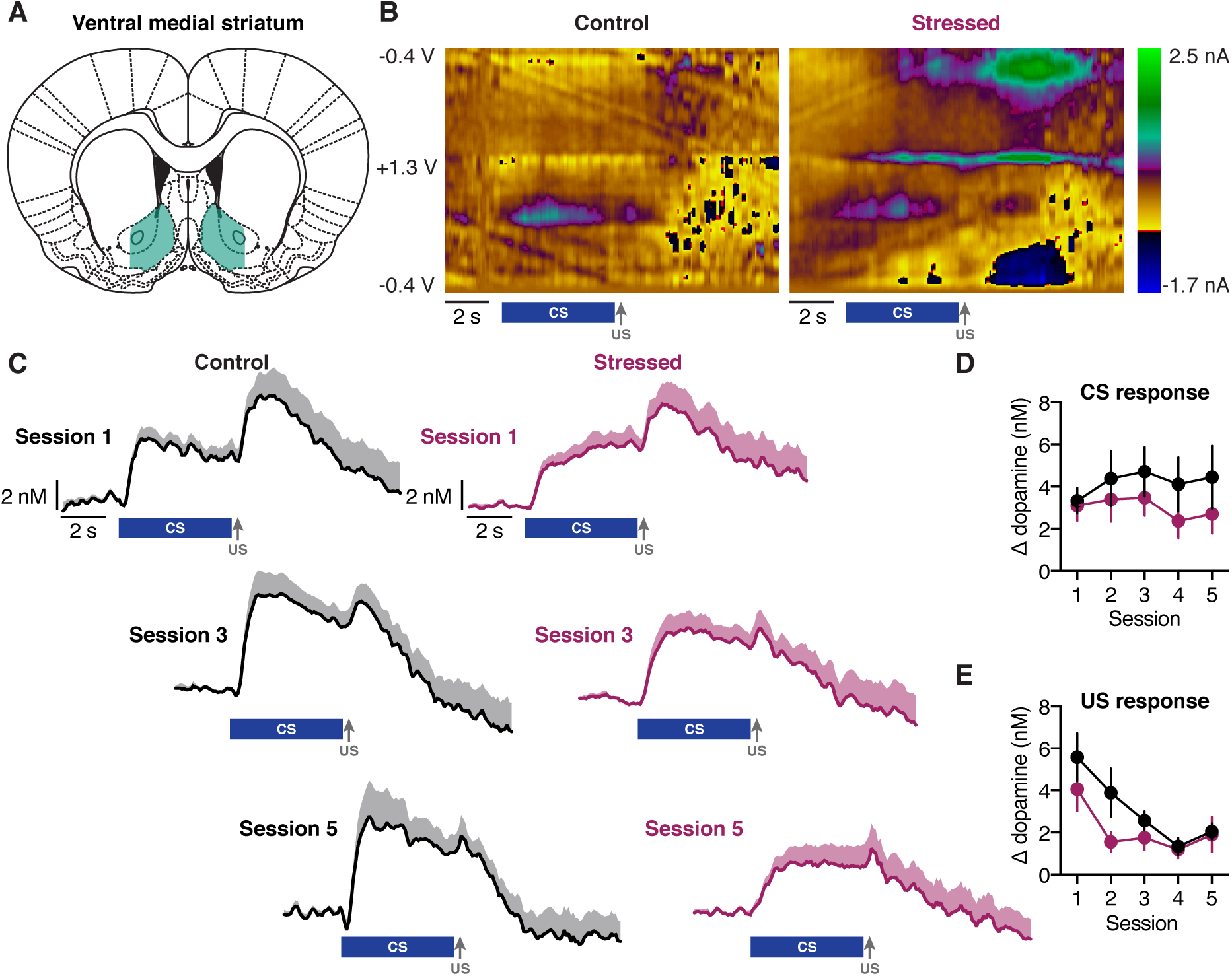
Acute stress does not alter dopamine signals in the VMS. **A**, Voltammetry recordings were taken from the VMS (shaded in cyan). **B**, Representative color plots of voltammetry recording during session 3 from a single electrode in a control rat (left) and a stressed rat (right). **C**, Average dopamine signals across electrodes in control rats (left) and stressed rats (right) during the first, third, and fifth training sessions. The blue bar denotes CS presentation and the grey arrow denotes reward delivery. **D**, Average CS-evoked dopamine release. **E**, Average US-evoked dopamine release.

Recent studies have illustrated heterogeneity of VMS dopamine responses to rewarding stimuli along the dorsal-ventral axis^38,39^. We compared VMS dopamine responses during the first training session from electrodes dorsal and ventral to the anterior commissure. There was no difference in CS- or US-evoked dopamine release between dorsal and ventral electrodes in control rats (CS: unpaired *t*-test *t*_(5.9)_=1.65, *p*=0.15; US: *t*_(5.0)_=0.28, *p*=0.79; *n=*5 dorsal, 5 ventral) or stressed rats (CS: *t*_(4.2)_=1.15, *p*=0.31; US: *t*_(8.5)_=0.0072, *p*=0.99; *n=*6 dorsal, 5 ventral).

Increasing evidence highlights that the ventral lateral striatum (VLS) contributes to reward-related behaviors^35,40,41^. Furthermore, aversive experience increases excitatory transmission to dopamine neurons projecting to the VLS^42^. As such, acute stress could enhance dopamine signaling in the VLS. Similar to the VMS, we identified time-locked dopamine signals in the VLS in response to the CS and US across Pavlovian conditioning sessions (Fig. 4B-C). There was no difference in CS-evoked dopamine release between stressed and control rats (treatment effect *F*_(1, 21)_=0.92, *p*=0.35, *n*=12 control, 13 stress; Fig. 4D). VLS dopamine release to the US decayed with training (session effect *F*_(2.6, 46.2)_=5.30, *p*=0.005; Fig.4E). However, US-evoked dopamine release was elevated across sessions in stressed animals (treatment effect *F*_(1, 21)_=8.16, *p*=0.01). Stress therefore selectively upshifts reward-evoked dopamine signals in the VLS.

**Figure 4.**
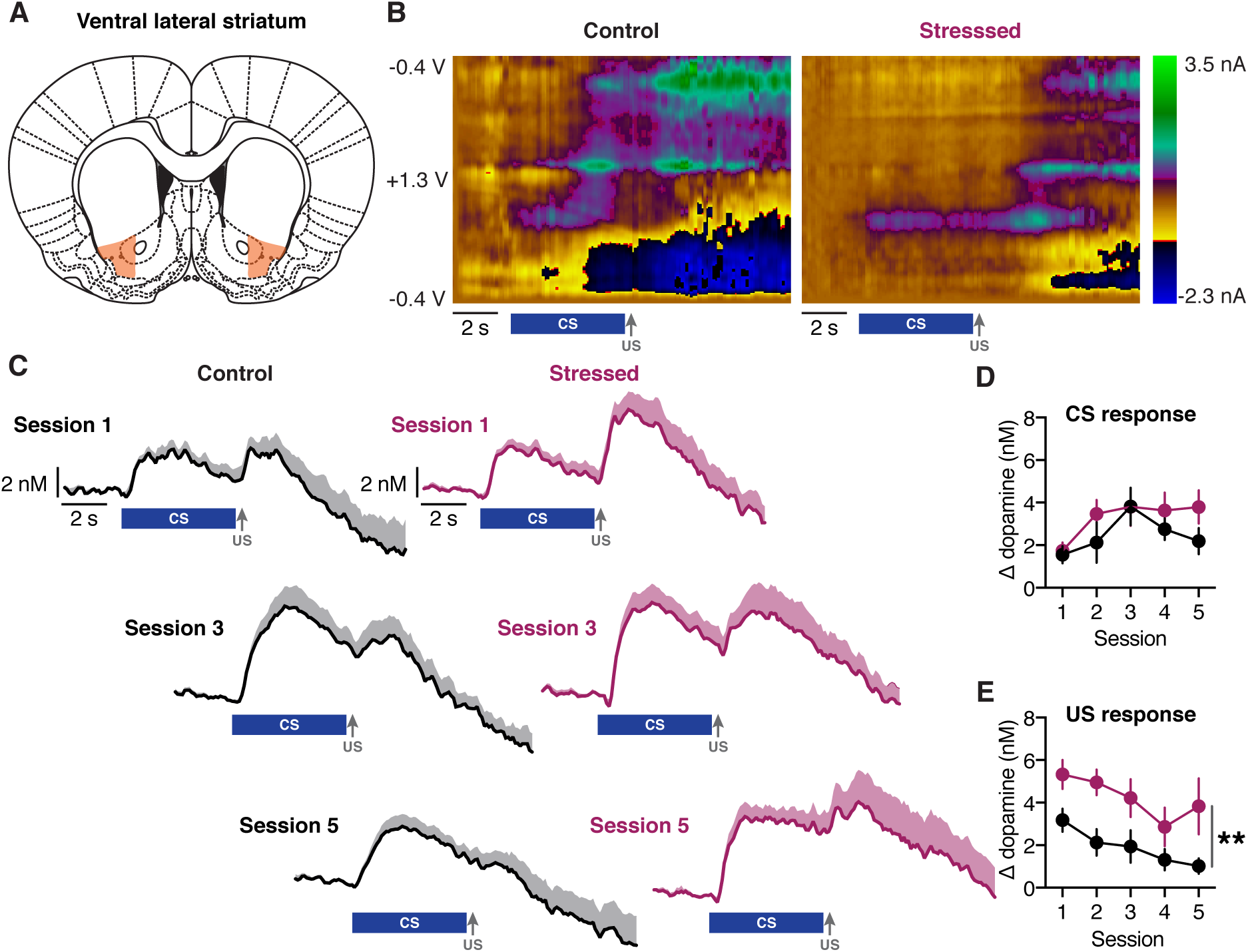
Acute stress selectively enhances US-evoked dopamine signals in the VLS. **A**, Voltammetry recordings were taken from the VLS (shaded in orange). **B**, Representative color plots of voltammetry recording during session 3 from a single electrode in a control rat (left) and a stressed rat (right). **C**, Average dopamine responses across electrodes in control rats (left) and stressed rats (right) during the first, third, and fifth training sessions. The blue bar denotes CS presentation and the grey arrow denotes reward delivery. **D**, Average CS-evoked dopamine release does not differ between groups. **E**, Average US-evoked dopamine release is enhanced in stressed rats. \*\**p* <. 01

### Stress recruits VLS dopamine signaling to regulate appetitive learning

We next examined whether VLS dopamine signaling was required for the stress-induced enhancement of conditioned responding. To address this, rats were implanted with bilateral cannulae targeting the VLS for local pharmacological manipulations. The D1/D2 dopamine receptor antagonist flupenthixol (10 µg/side) or vehicle was infused into the VLS 30 min before the first 5 sessions. Rats were trained without injections for 5 additional sessions to differentiate the acute versus sustained behavioral effects of the flupenthixol treatment (Fig. 5A).

**Figure 5.**
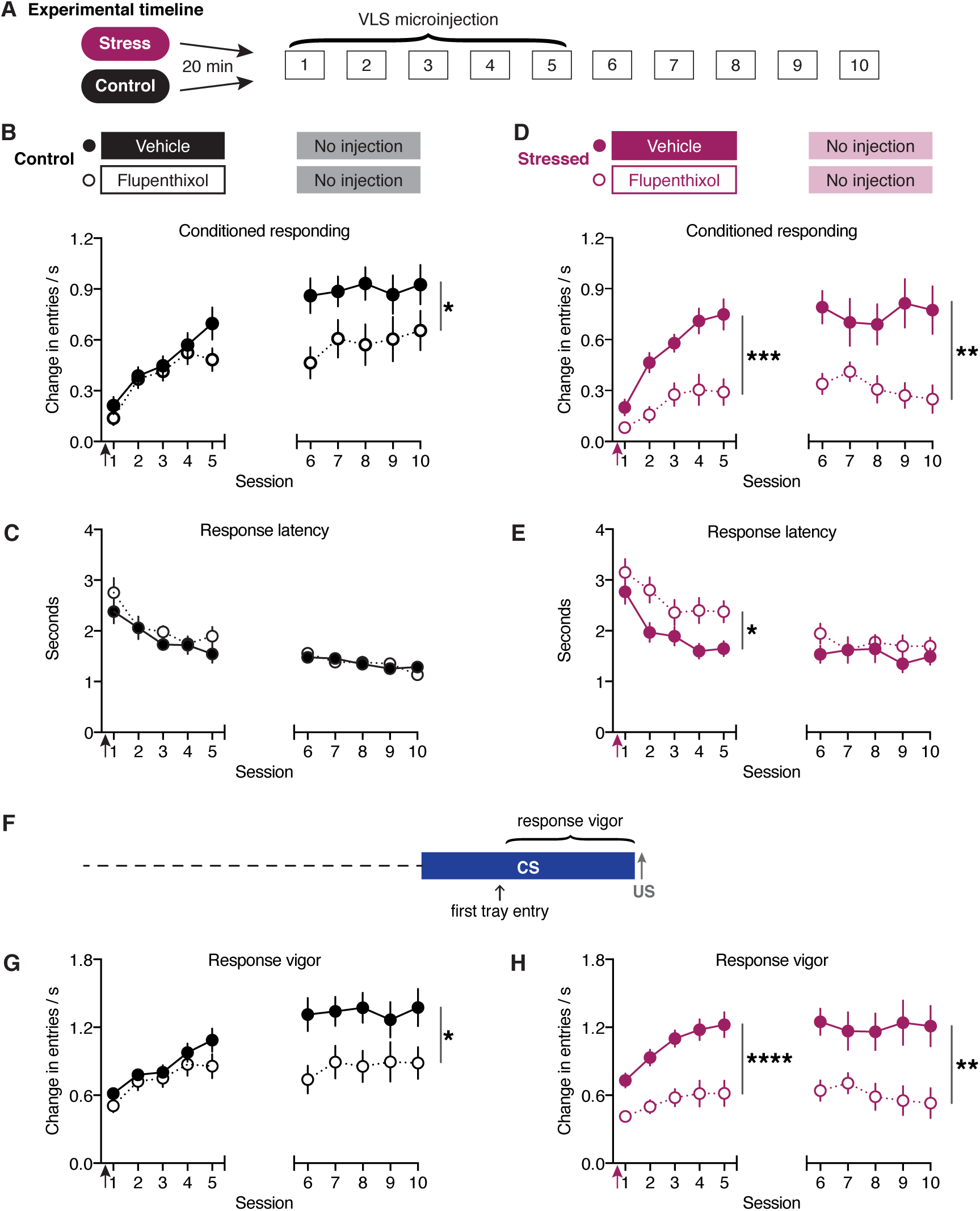
VLS dopamine signals are required for conditioning in stressed animals. **A**, Training paradigm. Animals were stressed once, 20 min prior to the first session. Flupenthixol or vehicle was infused to the VLS before each of the first 5 training sessions. Training continued for 5 additional sessions without injections. **B**, Conditioned responding acquisition was not initially impaired by flupenthixol treatment in control rats, but a delayed deficit emerged with additional training. **C**, Response latency was not altered by flupenthixol treatment in controls. **D**, Flupenthixol treatment impaired conditioned responding in stressed rats. **E**, Flupenthixol treatment reversibly increased the latency in stressed rats. **F**, Calculation of response vigor. **G**, Response vigor was not initially impaired by flupenthixol treatment in control rats, but a delayed deficit emerged with extended training. **H**, Flupenthixol treatment impairs response vigor in stressed rats. \**p* < .05, \*\**p* < .01, \*\*\**p* < .001, \*\*\*\**p* < .0001

VLS dopamine receptor blockade did not disrupt conditioned responding in control rats (drug effect *F*_(1, 20)_=1.45, *p*=0.24, *n*=11 vehicle, 11 flupenthixol; Fig. 5B). We note a non-significant reduction in responding on the last day of flupenthixol treatment and a significant reduction in subsequent sessions in which no injection was administered (prior treatment effect *F*_(1, 20)_=6.03, *p*=0.02; Fig. 5B). This suggests that VLS dopamine signaling is not essential for conditioned responding early in training but regulates behavior in later sessions. In contrast to the effect on conditioned responding, flupenthixol did not affect response latency acutely (drug effect *F*_(1, 20)_=1.37, *p*=0.26) or during subsequent sessions without injections (prior treatment effect *F*_(1, 20)_=0.0006, *p*=0.98; Fig. 5C). VLS dopamine receptor antagonism therefore selectively affects conditioned responding without altering approach latency in unstressed rats.

In stressed rats, flupenthixol in the VLS acutely suppressed conditioned responding (drug effect *F*_(1, 21)_=19.93, *p*=0.0002, *n*=12 vehicle, 11 flupenthixol; Fig. 5D). This effect persisted throughout subsequent sessions in which no injection was administered (prior treatment effect *F*_(1, 21)_=10.64, *p*=0.004; Fig. 5D). Furthermore, flupenthixol acutely slowed the latency to respond in stressed rats (drug effect *F*_(1, 21)_=6.41, *p*=0.02), but this effect was not observed in subsequent sessions (prior treatment effect *F*_(1, 21)_=0.92, *p*=0.35; Fig. 5E). Collectively, these results demonstrate that flupenthixol impaired conditioned responding earlier in training in stressed animals relative to controls. Furthermore, flupenthixol treatment reversibly increased response latency in stressed animals but had no effect in controls. These results highlight that conditioned responding and response latency are differentially regulated by VLS dopamine transmission.

The effect of flupenthixol treatment on conditioned responding in stressed animals could be driven by the increased latency to approach the food tray, which reduces the time available for conditioned head entries. To eliminate the confound of response latency, we recalculated the CS-evoked response rate based on the interval between the first head entry and the US delivery (‘response vigor’; Fig. 5F). In stressed rats, flupenthixol acutely reduced response vigor (drug effect *F*_(1, 21)_=33.80, *p* <0.0001; Fig. 5H), and this effect persisted throughout subsequent sessions (prior treatment effect *F*_(1, 21)_=10.37, *p* =0.004). These results illustrate that VLS dopamine transmission regulates both the latency and the vigor of conditioned appetitive responses in stressed animals.

## Discussion

In adverse circumstances, it is adaptive to rapidly and effectively learn which stimuli predict beneficial outcomes. Prior rodent studies have shown that stress enhances the learned preference for a cocaine-associated context^13,14^, though it was unclear if acute stress similarly facilitated learning driven by natural rewards. Here, we addressed this question by utilizing a Pavlovian task in which an auditory CS signaled the upcoming delivery of a food reward. We demonstrate that a single, brief episode of restraint stress induces a persistent increase in conditioned responding.

The effect of stress on subsequent behavior depends on the time elapsed from the stressor, as well as the duration, intensity, and frequency of the stressful experience^43-45^. Our results indicate that the influence of acute restraint stress on reward learning is time-dependent. Stress administered two hours prior to the first conditioning session failed to affect behavior. Additionally, acute stress did not increase conditioned responding in rats that had already learned the task. Stressful experience therefore has maximal influence over behavior when it occurs early in training. Collectively, these findings demonstrate that stress induces a short-term state that interacts with the associative learning mechanism to produce a long-term change in behavior.

Studies examining the role of ventral striatal dopamine in appetitive behavior have primarily focused on the VMS^26,37,46-49^. However, recent findings indicate that dopaminoceptive VLS spiny projection neurons regulate aspects of reward-seeking^35,40,41^. Our results demonstrate that dopamine in the VLS regulates conditioned responding in later training sessions in unstressed animals. Interestingly, acute stress shifts the temporal window in which VLS dopamine controls conditioned responding to an earlier point in training.

Stress selectively enhanced dopamine release in the VLS without affecting dopamine release in the VMS. This result is in line with previous findings demonstrating that the dopamine neurons targeting the VLS are anatomically and functionally distinct from those targeting the VMS^38,42,50-52^. Furthermore, VMS and VLS spiny projection neurons innervate different downstream targets, (e.g., medial vs. lateral ventral pallidum and VTA)^53,54^. Reward-evoked dopamine signals encode subjective value based upon one’s internal state (e.g., satiety)^55-57^. We suggest that the stress-induced increase in VLS dopamine release reflects an upshift in reward value which then invigorates conditioned appetitive behavior. Interestingly, our data illustrate that increased reward-evoked dopamine release accompanies invigorated CS-evoked behavior. We propose that the US-evoked dopamine signal initiates sustained changes downstream of the VLS, resulting in a persistent increase in conditioned responding.

A single traumatic experience can exert long-lasting effects on behavior, as is the case in post-traumatic stress disorder. As such, the role of stress in behavior motivated by aversive stimuli has been studied extensively. However, traumatic stress also alters responsivity to rewards^58,59^. Here, we demonstrate that a single stress exposure acts upon a specific mesolimbic circuit to produce lasting changes in appetitive behavior. These findings highlight the ventral lateral striatum as a nexus for stress to modulate the neural representation of reward.

## Acknowledgements

This work was supported by National Institutes of Health grants DA033386 and DA042362 to M.J.W. The authors thank Merridee Lefner for critical input on the manuscript. The authors declare no conflict of interest.

## Author Contributions

CES, SCT, and YR performed the experiments and analyzed the data. CES and MJW designed the experiments and wrote the manuscript.

